## Supplementary Table 1 for "Intrinsic timescales as an organizational principle of neural processing across the whole rhesus macaque brain" (from another version of this preprint: version 1, 2021-10-07)

**Supplementary Table 1. ROIs for Figure 1**

|  | **CHARM/SARM Level** | **Code** | **Area** |
| --- | --- | --- | --- |
| Murray | 3 | 25 | OFC |
|  | 3 | 3 | ACC |
|  | 2 | 50 | LPFC |
|  | 3 | 92 | S1 |
|  | 3 | 95 | S2 |
|  | 5 | 115 | LIP |
|  | 5 | 232 | MT |
| Fascianelli + Cirillo | 3 | 25 | PFo |
|  | 6 | 63, 67, 68 | DLPFC |
|  | Custom | Custom | PFp |
|  | 5 | 81 | PMd |
| Cavanagh | 6 | 63, 67, 68 | DLPFC |
|  | 6 | 53, 70, 74, 75 | VLPFC |
|  | 6 | 10, 62 | ACC |
| Maisson | 5 | 22 | vmPFC |
|  | 6 | 4 | pgACC |
|  | 6 | 5 | sgACC |
|  | Custom | Custom | dACC |
| Nougaret | 3 | 43 | Striatum |
|  | 5 | 91 | STN |
|  | 6 | 56 | GPe |
